# Delayed protein translocation protects mitochondria against toxic CAT-tailed proteins

**DOI:** 10.1101/2024.11.20.624519

**Authors:** Nils Bertram, Toshiaki Izawa, Felix Thoma, Serena Schwenkert, Nikola Wagener, Christof Osman, Walter Neupert, Dejana Mokranjac

## Abstract

Ribosome-associated quality control (RQC) protects cells against toxic effects of faulty polypeptides produced by stalled ribosomes. However, mitochondria are vulnerable to C-terminal alanyl-and-threonyl (CAT)-tailed proteins that are generated in this process and faulty nuclear-encoded mitochondrial proteins are handled by the recently discovered mitoRQC. Here, we performed a genome-wide screen in yeast to identify additional proteins involved in mitoRQC. We found that Pth2, a peptidyl-tRNA hydrolase in the mitochondrial outer membrane, influences aggregation of CAT-tailed proteins without majorly affecting the CAT-tailing process itself. Peptidyl-tRNA hydrolase activity is essential during this process, yet the activity of Pth2 can be substituted by another peptidyl-tRNA hydrolase, upon proper localization. Our data suggest that Pth2 acts through modulating protein translocation and that mitochondrial proteostasis network is relieved through increased access of CAT-tailed proteins to cytosolic chaperones. Other hits obtained in the screen show that, in general, delayed protein translocation protects mitochondria against toxic CAT-tailed proteins.

## Introduction

Defective proteins are constantly generated at a low rate during translation, due to for example mutations or chemical damage of the mRNA or the ribosome itself. These aberrant nascent chains can have cytotoxic properties and form protein aggregates, subsequently sequestering chaperones and inducing proteotoxic stress (Pilla, Schneider, and Bertolotti 2017; Hipp, Kasturi, and Hartl 2019). To prevent protein toxicity at an early stage, the ribosome-associated protein quality control (RQC) system monitors and degrades translation products that emerge from stalled ribosomes (Filbeck et al. 2022; Inada and Beckmann 2024; Iyer et al. 2023; Joazeiro 2019; Howard and Frost 2021). After dissociation of the stalled 80S ribosome(Pisareva et al. 2011; Shao, von der Malsburg, and Hegde 2013; Shoemaker, Eyler, and Green 2010), the 60S subunit, with the associated peptidyl-tRNA, is recognized by Rqc2 which then recruits E3 ubiquitin ligase Ltn1 to the complex (Defenouillère et al. 2013; Lyumkis et al. 2014; Shao et al. 2015; Shen et al. 2015). Ltn1 ubiquitylates nascent chains at the lysine residues present in the close proximity of the ribosomal exit tunnel (Lyumkis et al. 2014; Shao et al. 2015; Bengtson and Joazeiro 2010). Aided by Rqc1, together with Cdc48 and its cofactors, ubiquitylation leads to extraction of nascent chains and their subsequent degradation by the proteasome (Defenouillère et al. 2013; Brandman et al. 2012; Verma et al. 2013). In addition to its role in recruiting Ltn1, Rqc2 has the ability to noncanonically elongate nascent chains by adding C-terminal alanine and threonine residues in a template- and 40S- independent manner (Shen et al. 2015; Defenouillère et al. 2016; Tesina et al. 2023; Yonashiro et al. 2016; Osuna et al. 2017). This C-terminal modification of the nascent chain - termed CAT-tailing - is thought to facilitate ubiquitylation by Ltn1 by extruding additional lysine residues from the ribosomal exit tunnel (Kostova et al. 2017). CAT-tails were shown to serve as degrons in different organisms (Sitron and Brandman 2019; Lv et al. 2024; Lytvynenko et al. 2019; Thrun et al. 2021), however, they also render proteins aggregation-prone, leading to formation of SDS-insoluble aggregates (Choe et al. 2016; Yonashiro et al. 2016; Izawa et al. 2017; Wu et al. 2019).

RQC acting on stalled ribosomes that are engaged in synthesis of nuclear-encoded mitochondrial proteins is complicated by the presence of the N-terminal mitochondrial targeting sequences (MTS) (Izawa et al. 2017). The MTS namely can already engage with mitochondrial protein translocases, the TOM complex in the outer and the TIM23 complex in the mitochondrial inner membrane, and initiate translocation into the organelle, before the nascent chain is released from the 60S subunit. This results in tight apposition of the 60S subunit with the TOM complex which reduces access of Ltn1 to the nascent chains, hindering their ubiquitylation and subsequent extraction and degradation in the cytosol. However, CAT-tailing can still occur, eventually resulting in import of CAT-tailed proteins into mitochondria (Lv et al. 2024; Izawa et al. 2017). Mitochondria were recently shown to be especially sensitive to the presence of CAT-tailed proteins within the organelle, leading to a collapse of the mitochondrial proteostasis network, compromising mitochondrial function and ultimately resulting in cell death (Izawa et al. 2017). Therefore, mitochondria rely on the protective function of the cytosolic endonuclease Vms1 (Heo et al. 2010) which acts as a safeguard by halting CAT-tailing, both through peptidyl-tRNA cleavage (Verma et al. 2018; Kuroha et al. 2018) and displacement of Rqc2 from 60S subunit (Izawa et al. 2017; Su et al. 2019), which results in the release of aberrant but not CAT-tailed proteins into mitochondria. This recently identified RQC pathway acting on the mitochondrial surface is termed the mitoRQC (Izawa et al. 2017).

To identify additional proteins involved in mitoRQC, we conducted genome-wide screens in yeast. We identified Pth2, a functionally poorly characterized peptidyl-tRNA hydrolase in the mitochondrial outer membrane (De Pereda et al. 2004; Rosas-Sandoval et al. 2002; Schulte et al. 2023), to be a release factor of CAT-tailed proteins arrested in the TOM complex. Data presented here indicate that Pth2 does not influence the CAT-tailing of nascent chains but rather their translocation into mitochondria, providing more time for the cytosolic proteostasis network to deal with the mitoRQC substrates. Deletions of nonessential components of the TOM and TIM23 complexes had similar effects, suggesting that delaying protein translocation is a general mechanism protecting mitochondria against toxic CAT-tailed proteins.

## RESULTS AND DISCUSSION

### Pth2 is involved in mitoRQC

To identify novel proteins involved in mitoRQC, we employed unbiased, genome-wide screens in yeast *Saccharomyces cerevisiae* looking for components that would be able to rescue the growth defect of *Δvms1Δltn1* cells (Izawa et al. 2017) and thus produce strains viable on non-fermentable medium at 37°C. The first approach, in which we searched for high-copy suppressors, however, did not identify any obvious novel candidates, though Ltn1 and Vms1 were both repeatedly identified (data not shown). In the second approach, we mated the *Δvms1Δltn1* double mutant with the yeast deletion library and screened for the triple mutants that are viable under the screening conditions (Figure 1A). Rqc2 was present among the identified candidates, confirming the validity of the approach (Table S1). One candidate, peptidyl-tRNA hydrolase 2 (Pth2), drew our attention as it was previously shown to be an outer mitochondrial membrane protein which interacts with the TOM complex (Schulte et al. 2023) and its human homologue, PTRH2, was implicated in a number of human disorders (Sharkia et al. 2023). To validate the results obtained in the screen, we deleted *PTH2* in wild type, *Δvms1* and *Δvms1Δltn1* strains and conducted comprehensive growth analyses using serial drop dilution assay on both fermentable (YPD) and non-fermentable (YPG) media. Like in the screen, additional deletion of *PTH2* partially rescued the growth defect of *Δvms1Δltn1* on both fermentable and nonfermentable media, though not to the same extent as deletion of *RQC2* did (Figure 1B). We observed no growth defect of any of the single deletion strains nor of the *Δvms1Δpth2* double mutant under any of the conditions tested (Figures 1B and S1A). The growth phenotypes observed on solid media were also recapitulated in liquid cultures (Figure S1B).

**Figure 1:**
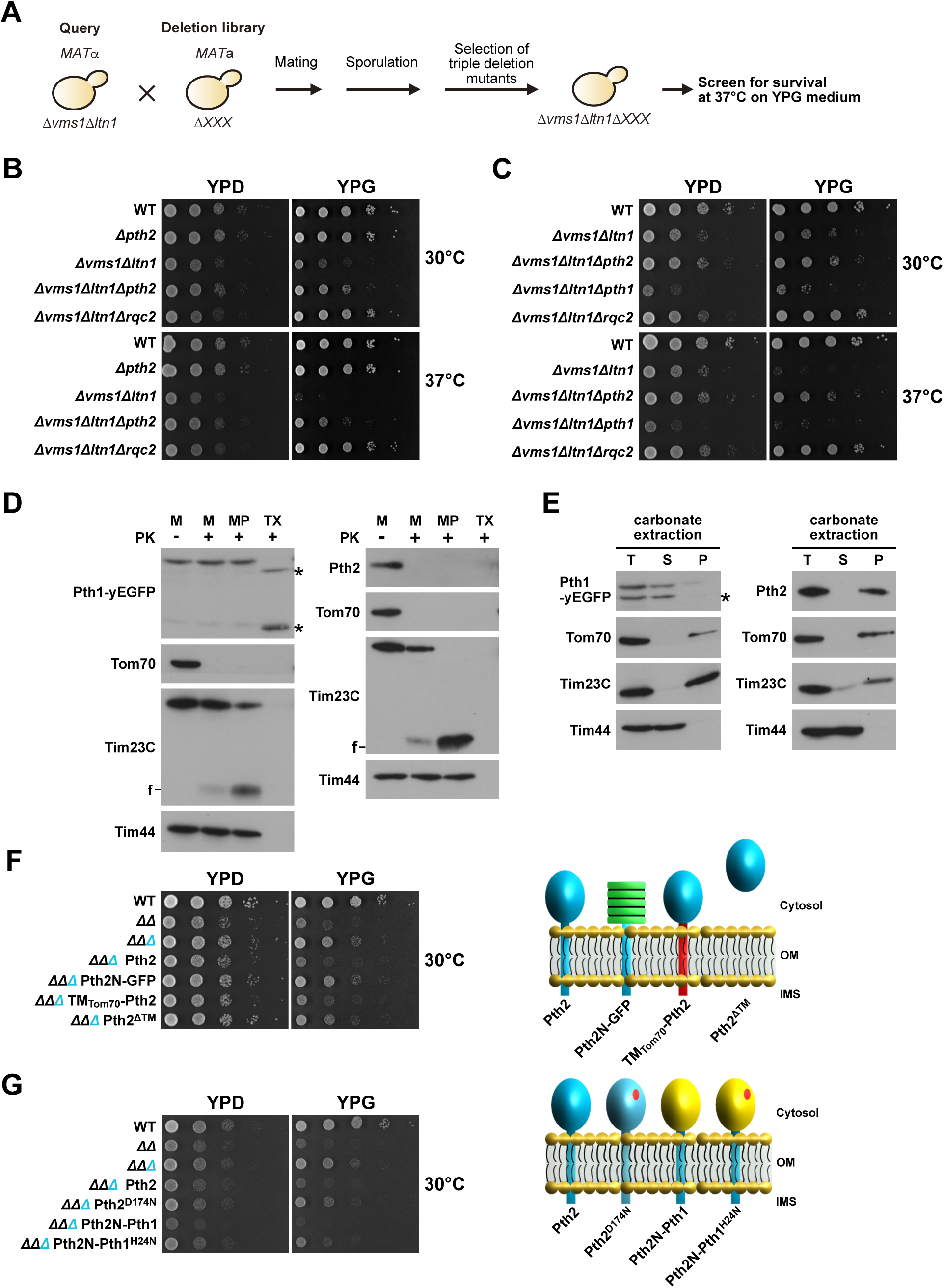
Pth2 is a peptidyl-tRNA hydrolase involved in the yeast mitoRQC. [A] Schematic description of the Systematic Genetic Array analysis. *Δvms1Δltn1* double deletion strain was crossed with a complete yeast deletion library. After sporulation, haploid triple deletion strains were tested for growth on a non-fermentable carbon source (YPG) at 37°C. [B] Growth of the indicated yeast strains was analyzed by a serial dilution assay. Serial 10-fold dilutios of cells grown in fermentable media in logarithmic phase were prepared and spotted on plates containing a fermentable (YPD) or a non-fermentable carbon source (YPG). Plates were incubated at 30°C or 37°C. [C] Growth of indicated yeast strains was analyzed as in 1B. [D] Isolated mitochondria (M), mitoplasts (MP) prepared by osmotic shock and Triton-solubilized mitochondria (TX) were treated with Proteinase K (PK), where indicated. Samples were analyzed by SDS-PAGE and western blotting with the indicated antibodies. Tom70, Tim23 and Tim44 were used as markers for the outer mitochondrial membrane, inner mitochondrial membrane and the mitochondrial matrix, respectively. f - stable fragment of Tim23 in the inner membrane; * - degradation products of Pth1-yEGFP. Pth1-yEGFP was detected using antibodies against GFP. [E] Total (T), supernatant (S) and pellet (P) fractions of carbonate extraction were analyzed by SDS-PAGE and western blotting using indicated antibodies. [F, G] Growth of the indicated yeast strains was analyzed as described in Fig 1B (left panels). Cartoon representations of the analyzed Pth2 constructs (right panels). *ΔΔ = Δvms1Δltn1; ΔΔΔ = Δvms1Δltn1Δpth2*

In addition to Pth2, mitochondria contain another peptidyl-tRNA hydrolase, Pth1 (Rosas-Sandoval et al. 2002), whose mammalian homologue, Ptrh1, was recently implicated in the RQC pathway in mammalian cells (Kuroha et al. 2018). In addition, bacterial Pth was recently shown to release stalled nascent chains in a CAT-tail dependent manner (Svetlov et al. 2024). Interestingly, we did not identify Pth1 in the screen. To exclude potential artefacts of the high-throughput screening, we deleted *PTH1* in both wild type and *Δvms1Δltn1* yeast strains. In contrast to the deletion of *PTH2*, deletion of *PTH1* did not improve growth of *Δvms1Δltn1* cells but rather aggravated the already severe growth defect (Figures 1C and S1C). This observation can be explained by the different subcellular localizations of the two enzymes, supporting their involvements in different pathways. Pth1 is a soluble protein in the mitochondrial matrix and Pth2 is integrated in the outer membrane and exposing its catalytic domain to the cytosol (Figures 1D and 1E). Whether Ptrh1 in mammalian cells may localize to the mitochondrial surface, under normal or stress conditions, remains to be determined. In conclusion, Pth2 and not Pth1 is involved in the mitoRQC in yeast.

Based on the evolutionary conservation of its primary sequence, Pth2 was previously divided into N- and C-terminal regions (De Pereda et al. 2004; Ishii, Funakoshi, and Kobayashi 2006) (Figure S1D). While the N-terminal region lacks conservation on the primary sequence level, it is predicted to contain a transmembrane domain in all examined orthologues (Figures S1D and S1E). The C-terminal region, on the other hand, is highly conserved among different eukaryotic species (Figure S1D). To investigate the requirement for the N- and the C-terminal regions of Pth2 within the mitoRQC, we generated different Pth2 constructs (Figures 1F, 1G and S1F). The wild type Pth2 construct, expressed under the control of the *PTH2* promoter and genomically integrated into the *LEU* locus of *Δvms1Δltn1Δpth2* cells, successfully reconstituted the *Δvms1Δltn1* phenotype (Figures 1F, 1G, S1F and S1G), validating the approach taken. The substitution of the entire C-terminal region of Pth2 with superfolded GFP (Pth2N-GFP) led to a strain that phenocopied *Δvms1Δltn1Δpth2*, suggesting that this construct is a functionally inactive form of Pth2 (Figures 1F, S1F and S1G). In contrast, the replacement of the transmembrane domain of Pth2 with that of the outer mitochondrial membrane protein Tom70 (TM_Tom70_-Pth2), to maintain the localization of the C-terminal region of the protein, replicated the effect of the expression of wild type Pth2, suggesting that the C-terminal region of Pth2 plays an essential role in the mitoRQC (Figures 1F, S1F and S1G). The construct lacking the transmembrane domain (Pth2^ΔTM^), which was previously shown to reside in the cytosol (Schulte et al. 2023), showed an intermediate phenotype (Figure 1F, S1F, S1G), indicating that the mitochondrial localization supports the function of Pth2 within the mitoRQC.

To assess whether the catalytic activity of Pth2 is important within the mitoRQC, we introduced a point mutation (Pth2^D174N^) in its catalytic site that was previously demonstrated to impair the peptidyl-tRNA hydrolase activity of Pth2 (Ishii, Funakoshi, and Kobayashi 2006). Cells expressing Pth2^D174N^ in *Δvms1Δltn1Δpth2* background grew better than the *Δvms1Δltn1* cells (Figures 1G, S1F and S1I), strongly suggesting that the peptidyl-tRNA hydrolase activity of Pth2 is required within the mitoRQC. We wondered whether this function is exclusive for Pth2 or if another peptidyl-tRNA hydrolase could substitute for it. To this end, we fused Pth1 to the C-terminus of the N-terminal region of Pth2, to ensure localization of Pth1 in the outer membrane (Figures 1G, S1F and S1I). Expression of the Pth2N-Pth1 fusion construct in the *Δvms1Δltn1Δpth2* background impaired growth of yeast cells even stronger than the *Δvms1Δltn1* strain has (Figures 1G and S1I). However, introduction of a point mutation that abrogates the catalytic activity of Pth1 (Pth2N-Pth1^H24N^) (Kuroha et al. 2018), improved cell growth (Figures 1G, S1F, S1H and S1I).

Taken together, these data indicate that Pth2 is involved in the mitoRQC pathway as a peptidyl-tRNA hydrolase. The function of Pth2 can, however, be taken over by another peptidyl-tRNA hydrolase, as long as it is correctly localized in the cell. This finding is in agreement with previous work which demonstrated a remarkable functional conservation of peptidyl-tRNA hydrolyses - both mitochondrial Pths can substitute for the essential enzyme in bacteria (Ishii, Funakoshi, and Kobayashi 2006; Rosas-Sandoval et al. 2002).

### Absence of Pth2 reduces aggregate formation but does not affect CAT-tailing

The toxicity associated with the dysfunction of the mitoRQC in *Δvms1Δltn1* cells is primarily attributed to CAT-tail-dependent aggregation of mitochondrial proteins within the mitochondrial matrix (Izawa et al. 2017). To investigate if the absence of Pth2 mitigates mitochondrial aggregate formation, we isolated detergent-insoluble protein aggregates from cell lysates of WT, *Δvms1Δltn1*, *Δvms1Δltn1Δpth2*, *Δvms1Δltn1Δrqc2* and *Δvms1Δltn1Δpth2* cells with reintroduced Pth2. Rieske Fe-S protein, Rip1, which was previously used as a marker for aggregation of mitochondrial proteins (Izawa et al. 2017), was present in the soluble fraction and was completely processed to its mature form in WT cells but was exclusively found in the aggregate fraction and was processed only to its intermediate form in cell lysates of *Δvms1Δltn1* cells (Figure 2A), as described previously (Izawa et al. 2017). Whereas additional deletion of *RQC2*, in *Δvms1Δltn1Δrqc2* cells, fully restored both the solubility and processing of Rip1, in *Δvms1Δltn1Δpth2* cells only part of Rip1 was fully processed and soluble (Figure 2A). A fraction of the intermediate form of Rip1 was, however, also present in the soluble form. In *Δvms1Δltn1Δpth2* cells with reintroduced Pth2, Rip1 behaved essentially the same as in *Δvms1Δltn1.* Ssq1, an Hsp70 chaperone in the mitochondrial matrix, behaved similarly to Rip1 in all cell lysates. In contrast, the cytosolic protein Pgk1 was only present in the soluble fraction. Thus, additional deletion of *PTH2* partially improved solubility of mitochondrial proteins and restored mitochondrial proteostasis of *Δvms1Δltn1* cells, correlating well with the observed partial rescue of growth.

**Figure 2:**
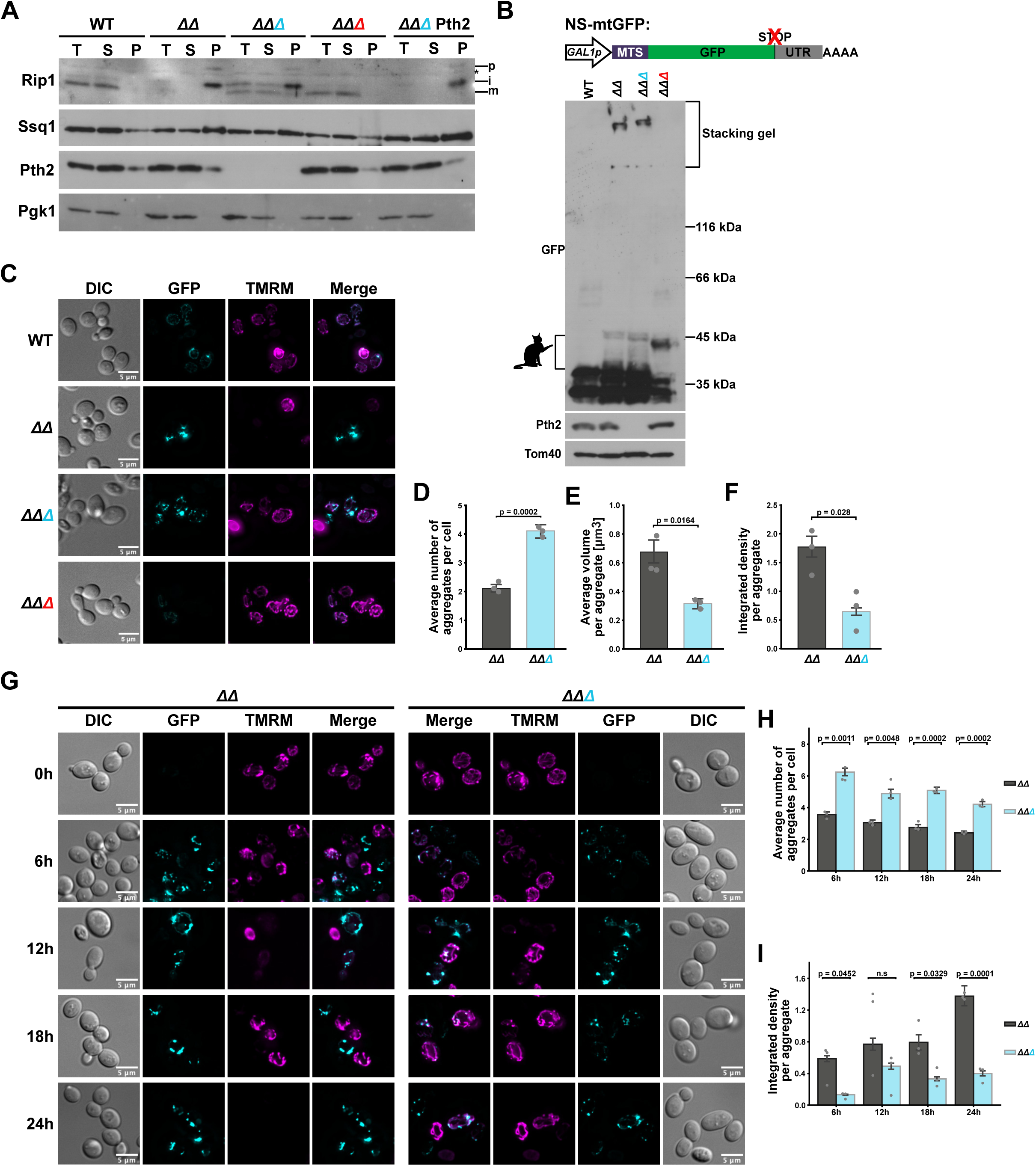
Absence of Pth2 reduces aggregate formation but does not affect CAT-tailing. [A] Indicated yeast cells were grown in YPGal medium at 37°C. Total cell extracts (T) were prepared and either directly analyzed by SDS-PAGE followed by western blot or first fractionated by centrifugation into supernatant fraction (S), containing soluble proteins, and pellet fraction (P), containing aggregated proteins. T and S, 10% of P. p, precursor, i, intermediate, and m, mature forms of Rip1; *, unspecific band. [B] Schematic representation of the NS-mtGFP construct used (upper panel). Expression of NS-mtGFP from a *GAL1* promoter was induced by addition of 0.5% galactose to cultures in logarithmic growth phase for 20h at 30°C. Total cell extracts were analyzed by SDS-PAGE and western blot. Stacking gel (Bracket) contains aggregated NS-mtGFP. Cat represents CAT-tailed forms of NS-mtGFP. [C] Fluorescence microscopy of NS-mtGFP-expressing cells. Yeast cells were grown as in 2B. Mitochondria were stained with a membrane potential sensitive dye TMRM. GFP signal was scaled equal, TMRM signal was auto-adjusted. [D, E, F] Aggregate number, volumes and integrated density (product of area and mean grey value) of microscopy images of *ΔΔ* and *ΔΔΔ* cells from 2C. The GFP channel of single cells was analyzed using the Fiji 3D object counter. Statistical analysis was done comparing the mean values of three biological replicates using a two-sided t-test. At least 150 cells per strain were analyzed in each replicate. Results are represented as mean with 95% confidence interval. [G] Fluorescence microscopy of *ΔΔ* and *ΔΔΔ* cells at different time points after induction of NS-mtGFP expression. Yeast cells growing in selective lactate medium in logarithmic growth phase were diluted to OD_600_ of 0.3 (0h). NS-mtGFP expression was subsequently induced by addition of galactose to 0.5%. After indicated time periods, samples were taken. Channels were adjusted as in 2C. [H-I] Quantification of aggregate number [H] and integrated density of aggregates [I] was done as in Figures 2D/F. *ΔΔ = Δvms1Δltn1; ΔΔΔ = Δvms1Δltn1Δpth2; ΔΔΔ = Δvms1Δltn1Δrqc2*

We next analyzed whether Pth2 affected CAT-tailing. To this end, we expressed a mitochondria-targeted version of GFP lacking the stop codon (NS-mtGFP) from an inducible *GAL1* promoter (Figure 2B). Translation of mRNAs lacking the stop codon results in proteins that contain a C-terminal poly-lysine stretch due to a partial translation of the mRNA polyA tails. These poly-lysine stretches stall the non-stop (NS) proteins in the ribosome exit tunnel making them substrates of RQC (Joazeiro 2017). CAT-tailing of this construct, visible as a smear running slower than the monomeric GFP (Brandman et al. 2012; Choe et al. 2016; Izawa et al. 2017), was obvious in total cell extracts of the *Δvms1Δltn1* and *Δvms1Δltn1Δpth2* strains. Furthermore, high molecular weight, SDS-insoluble aggregates were detected in the stacking gel of the same samples (Figure 2B). Neither CAT-tailing nor aggregation of GFP was detected in WT and *Δvms1Δltn1Δrqc2* cells. Therefore, Pth2 does not appear to majorly influence the CAT-tailing process in *Δvms1Δltn1* cells.

To further investigate protein aggregates present in *Δvms1Δltn1* and *Δvms1Δltn1Δpth2* cells, we used widefield fluorescence microscopy. Twenty hours after induction of NS-mtGFP expression, in WT and *Δvms1Δltn1Δrqc2* cells only a faint GFP signal was detected, which colocalized with mitochondrial tubules (Figures 2C and S2A). In contrast, *Δvms1Δltn1* and *Δvms1Δltn1Δpth2* cells clearly contained aggregated GFP material. However, these appeared smaller and less intense in *Δvms1Δltn1Δpth2* cells, compared to aggregates visible in *Δvms1Δltn1* cells (Figure 2C). Using a semi-automated analysis, we quantitatively analyzed number, size and intensity of the aggregates (Figures 2D-F), which confirmed our previous assessments. For additional visualization of mitochondria, we used membrane potential-sensitive dye TMRM (Chazotte 2011). While TMRM effectively stained mitochondria in the majority of WT and *Δvms1Δltn1Δrqc2* cells, we noticed that cells with large GFP aggregates showed no staining, suggesting a breakdown of mitochondrial membrane potential (not quantified). To assess the kinetics of aggregate formation in *Δvms1Δltn1* and *Δvms1Δltn1Δpth2* cells, we analyzed cells before and 6h, 12h, 18h and 24h after induction of NS-mtGFP expression. Whereas cells prior to induction had no visible GFP signal, aggregates were visible in both strains 6h post induction. The aggregates were notably smaller and less intense in *Δvms1Δltn1Δpth2* cells, compared to those in the *Δvms1Δltn1* cells (Figure 2G). Overall, *Δvms1Δltn1* cells showed a time-dependent decrease in number of aggregates per cell with a simultaneous increase in their intensity (Figure 2H and 2I). On the other hand, in *Δvms1Δltn1Δpth2* cells, the number of aggregates per cell exhibited only a minimal change across the different timepoints, accompanied with only a small increase in their intensity over time (Figure 2G-I).

In summary, deletion of *PTH2* reduces aggregate formation in *Δvms1Δltn1* cells but does not appear to majorly affect the CAT-tailing process.

### Absence of Pth2 delays import of CAT-tailed nascent chains by keeping them attached to the 60S subunit in the cytosol

Building upon our finding that Pth2 functions as a peptidyl-tRNA hydrolase within the mitoRQC, we hypothesized that it may play a role in the release of stalled polypeptide chains from the 60S subunit and that, in the absence of Pth2, the nascent chains may remain longer bound to 60S, which would effectively reduce their aggregation that we observed. To this end, we first analyzed whether NS-mtGFP-tRNA intermediates accumulated in the absence of Pth2. When cell lysates were prepared in the presence of an RNase inhibitor (DEPC), we observed an additional GFP band migrating at approximately 50kDa which was weak in the *Δvms1Δltn1Δpth2* cells and stronger in the *Δvms1Δltn1Δrqc2* cells (Figure 3A). Treatment with RNase A completely removed this species, confirming its identity as the peptidyl-tRNA (Verma et al. 2013). Considering the difference in the signal intensities of peptidyl-tRNA species in *Δvms1Δltn1Δpth2* and *Δvms1Δltn1Δrqc2* cells, it appears that, in the absence of both Vms1 and Pth2, another endonuclease and/or peptidyl-tRNA hydrolase can release nascent chains from the 60S. However, neither Pth2 nor this additional pathway seems to have access to non-CAT-tailed peptidyl-tRNAs which accumulated in the absence of Rqc2, Ltn1 and Vms1. Very strong accumulation of the peptidyl-tRNA species observed in *Δvms1Δltn1Δrqc2* cells may be the reason why deletion of *RQC2* did not restore growth of *Δvms1Δltn1* cells upon induction of NS-mtGFP expression (Figure S3A) – CAT-tailing may play a crucial role in the recycling of ribosomes in the presence of overwhelming amounts of RQC substrates.

**Figure 3:**
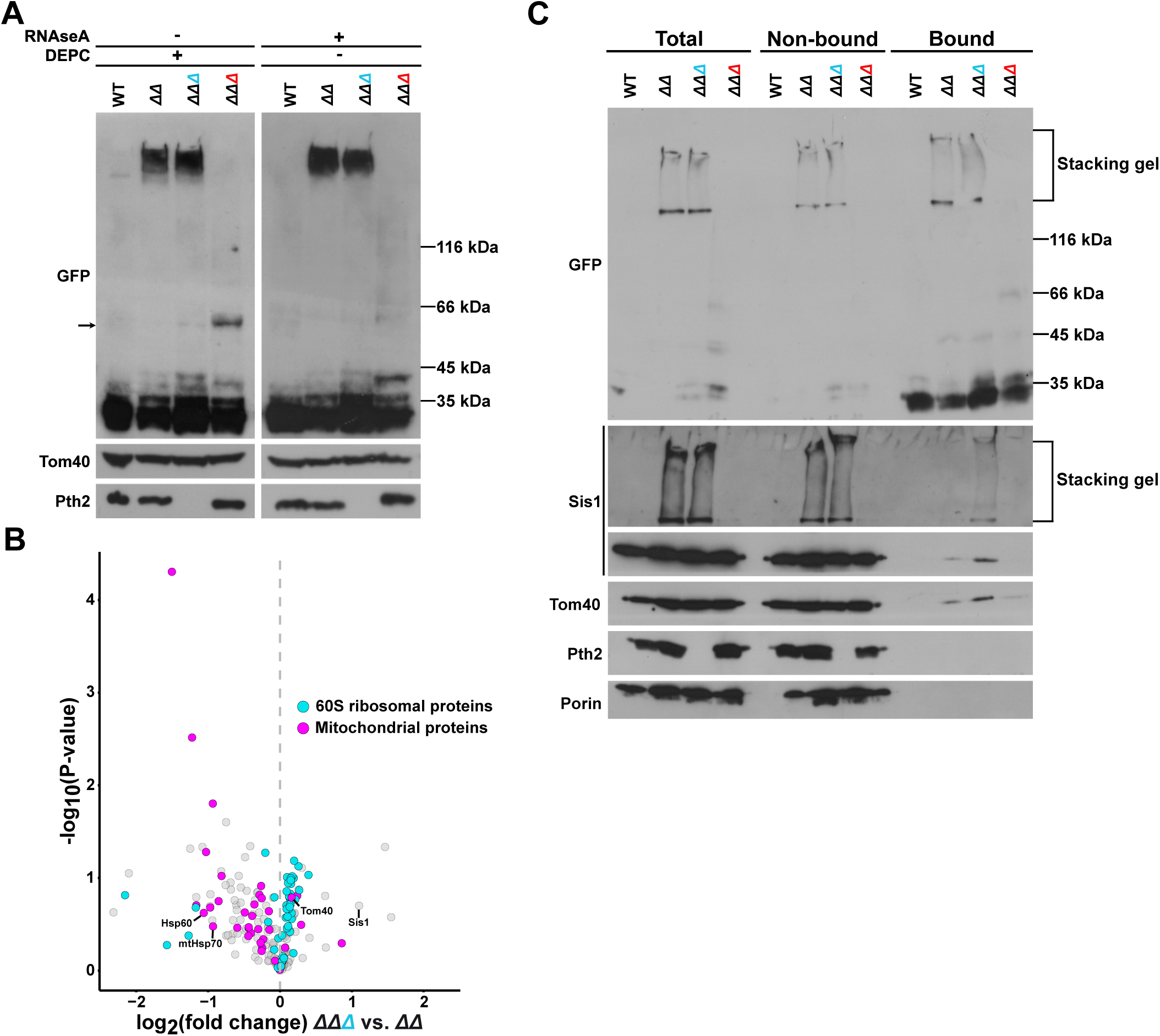
Absence of Pth2 reduces import of CAT-tailed nascent chains by keeping them attached to the 60S subunit in the cytosol. [A] Analysis of tRNA-linked NS-mtGFP. Cells were grown as in 2B and lysed either in presence of RNAseA or the RNAse inhibitor DEPC. Lysates were analyzed by neutral pH SDS-PAGE and western blot with the indicated antibodies. Arrow represents tRNA-NS-mtGFP conjugate. [B] Yeast cells were grown as in 2B. Lysates were subjected to immunoprecipitation with anti-GFP nanobodies followed by on-bead digestion and label-free quantification by mass spectrometry. Volcano plot of NS-mtGFP interactors in *ΔΔΔ* versus *ΔΔ* cells. Log_10_ P-value and log_2_ fold enrichment are shown. Protein constituents of the ribosomal 60S subunit are shown in blue and mitochondrial proteins in magenta. [C] Cell lysates as in 3B were immunoprecipitated using anti-GFP nanobodies Total, nonbound and bound fractions were analyzed by SDS-PAGE followed by immunoblotting and decoration with indicated antibodies. Total and nonbound fractions correspond to 10% of the material in bound fraction. *ΔΔ = Δvms1Δltn1; ΔΔΔ = Δvms1Δltn1Δpth2; ΔΔΔ = Δvms1Δltn1Δrqc2*

To obtain further insight into the role of Pth2 within mitoRQC, we analyzed the interactomes of NS-mtGFP in *Δvms1Δltn1Δpth2* and *Δvms1Δltn1* cells. For this, we immunoprecipitated NS-mtGFP and analyzed the samples using mass spectrometry (Figure 3B and Table S2). 60S ribosomal proteins clustered on the *Δvms1Δltn1Δpth2* side of the volcano plot, demonstrating a more stable interaction of NS-mtGFP with 60S subunit in *Δvms1Δltn1Δpth2* cells, compared to *Δvms1Δltn1* cells. This finding supports the notion of the delayed release of the stalled nascent chains from the 60S subunit in the absence of Pth2. Furthermore, the interactome of NS-mtGFP in *Δvms1Δltn1Δpth2* cells contained reduced levels of the majority of mitochondrial proteins, including the chaperones Hsp60 and mtHsp70 (Figures 3A-C), consistent with the decreased aggregation of mitochondrial proteins observed in these cells (Figure 2A-F). One notable exception was Tom40 which was enriched in the NS-mtGFP interactome in *Δvms1Δltn1Δpth2* cells (Figure 3B). The increased association of Tom40 with NS-mtGFP in *Δvms1Δltn1Δpth2* cells, compared to *Δvms1Δltn1* cells, was confirmed when the immunoprecipitates were analyzed by western blot (Figure 3C). Interestingly, one of the proteins with the highest enrichment in the interactome of NS-mtGFP in *Δvms1Δltn1Δpth2* cells was Sis1 (Figure 3B), the cytosolic chaperone previously shown to strongly interact with cytosolic CAT-tailed proteins (Choe et al. 2016; Defenouillère et al. 2016; Chang et al. 2024). We validated also this result of mass spectrometry using western blot (Figure 3C). Intriguingly, even though we observed aggregation of Sis1 in both *Δvms1Δltn1* and *Δvms1Δltn1Δpth2* cells, this aggregated Sis1 was specifically bound to NS-mtGFP only in *Δvms1Δltn1Δpth2* cells (Figure 3C).

### Delayed translocation protects mitochondria against toxic CAT-tailed proteins

The findings described above suggest that the NS-mtGFP-tRNA intermediate bound to 60S subunit is stabilized in the TOM pore in *Δvms1Δltn1Δpth2* cells, essentially preventing its import into mitochondria. We therefore wondered whether delayed translocation of CAT-tailed proteins could be a general mechanism used by cells to protect their mitochondria. To this end, we reanalyzed the list of candidates obtained in the screen. Indeed, virtually all nonessential subunits of the TOM and TIM23 complexes were present among the identified genes whose deletion restored growth of *Δvms1Δltn1* cells (Table S1). Here, we confirmed that additional deletions of *TOM6* from the TOM complex and of *TIM21* from the TIM23 complex rescued growth of *Δvms1Δltn1* cells essentially to WT levels, both on fermentable and nonfermentable carbon sources and both on 30°C and 37°C (Figure 4A and 4B).

**Figure 4:**
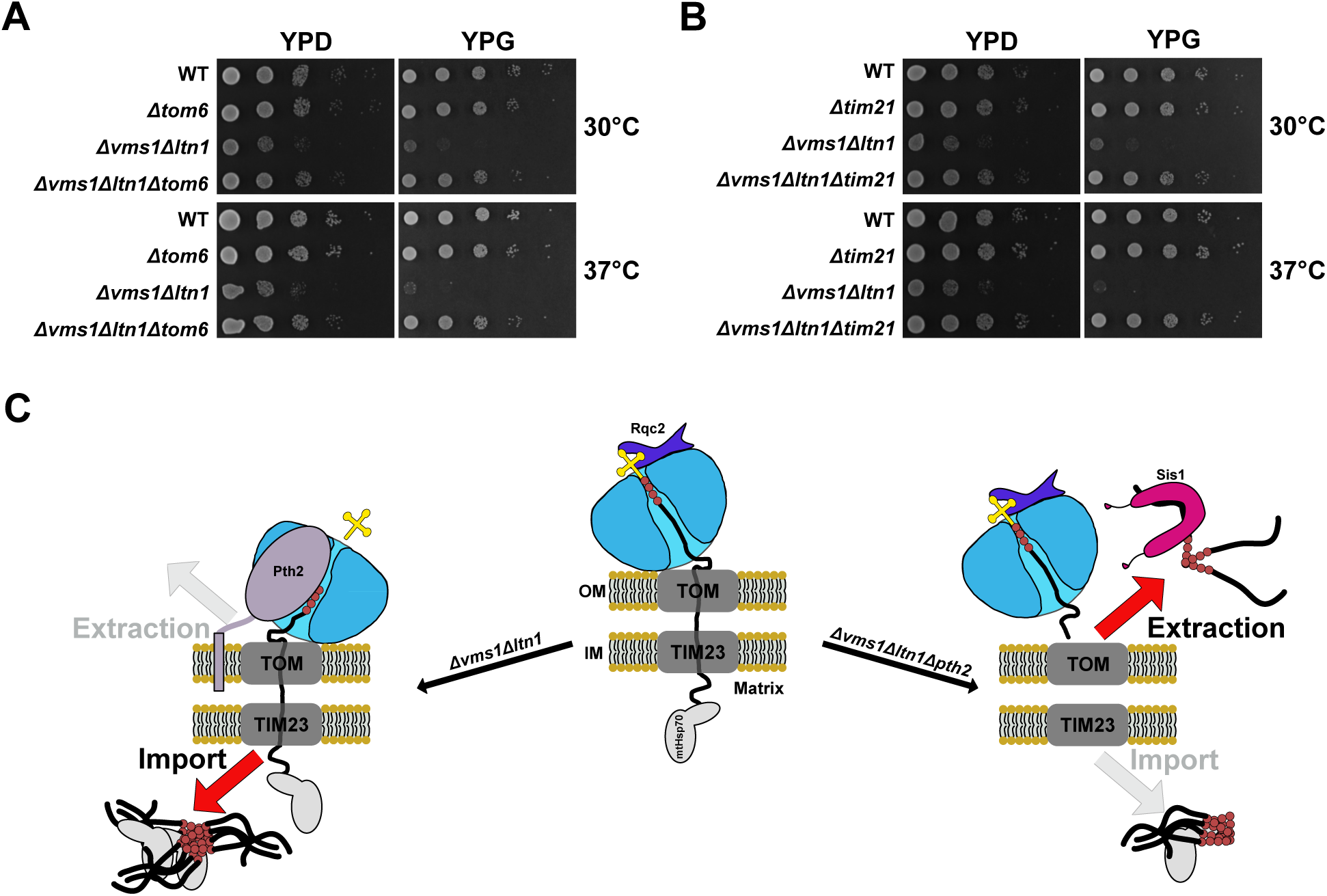
Delayed protein translocation protects mitochondria against toxic CAT-tailed proteins. [A, B] Growth of indicated yeast strains was analyzed as in 1B. [C] Model of Pth2 function within the mitoRQC. See text for details.

Based on the available data, we propose the following model of the role of Pth2 within mitoRQC (Figure 4C). Substrates of the mitoRQC are faulty nuclear-encoded mitochondrial proteins that are still bound to 60S and they accumulate in the TOM and TIM23 complexes on their way into mitochondria. In the absence of Vms1 and Ltn1, they are constantly CAT-tailed by Rqc2. These CAT-tailed nascent chains are substrates of the peptidyl-tRNA hydrolase Pth2 in the outer membrane. Pth2 releases them from the 60S, allowing their complete import into mitochondria. However, once in the mitochondrial matrix, these CAT-tailed proteins aggregate, sequestering mitochondrial chaperones and resulting in mitochondrial toxicity. In the absence of Pth2, release of CAT-tailed nascent chains from the 60S is delayed and they remain longer in the TOM complex, providing more time to the cytosolic proteostasis network and Sis1 in particular to act. The prolonged access of mitoRQC substrates to the cytosolic proteostasis network can also be achieved by lowering the activities of the TOM and TIM23 complexes. At the same time, prevention of import of CAT-tailed proteins into mitochondria relieves organellar proteostasis network and restores mitochondrial function.

## ACKNOWLEDGEMENTS

We would like to thank the current and former members of the Mokranjac Lab and the Mitoclub for their support and stimulating discussions. We also wish to thank Johannes Buchner for providing the antibody against Sis1 and Roland Beckmann for critical comments on the project. This research was generously supported by the Deutsche Forschungsgemeinschaft (projects NE101/28-1 to W.N. and MO1944/2-1 and MO1944/3-1 to D.M.). F.T. and C.O. were supported by a Human Frontier Science Program Research Grant (RGP021/2023) and a DFG Research Grant (OS 410/3-1).

## AUTHORS CONTRIBUTION

Conceptualization - D.M., W.N.; Methodology - D.M., N.B.; Formal Analysis - N.B., T.I., F.T., S.S.; Investigation - N.B., T.I., S.S., N.W.; Writing Original Draft - D.M., N.B.; Visualization - N.B., T.I., F.T.; Supervision - D.M., W.N., C.O.; Funding Acquisition - D.M., W.N., C.O.; All authors discussed the obtained results and commented on the original draft of the manuscript.

## ASUPPLEMENTAL INFORMATION

Document S1. Figures S1-3

Table S1. Results of the Synthetic Genetic Array analysis

Table S2. NS-mtGFP mass spectrometry interactome

Table S3. Strains, Plasmids and Primers used in the study

## MATERIAL AND METHODS

### Yeast strains and plasmids

The wild-type *Saccharomyces cerevisiae* strain BY4742 (*Mat* α *his3Δ1 leu2Δ0 lys2Δ0 ura3Δ0*) was used for all genetic manipulations. A comprehensive list of strains, primers and plasmids can be found in Table S3. Chromosomal tagging and deletions were made by homologous recombination of PCR products, as described previously (Janke et al. 2004). The junctions were all confirmed by PCR and, if possible, also on protein level.

The various Pth2 constructs were genomically integrated into the LEU locus in a markerless manner, as described in Schrott and Osman, 2023 (Schrott and Osman 2023). Briefly, DNA sequence encoding *PTH2*, together with ca. 500bp upstream and downstream of the ORF, was amplified from yeast genomic DNA and cloned into pSS036 (Schrott and Osman 2023) via Gibson assembly, exchanging the DNA sequence encoding mitochondrially targeted mKate2 under the control of the *PGK1* promoter and the *ADH1* terminator with the *PTH2* sequence along with its promoter and 3’UTR. This plasmid served as a template for generation of all other Pth2 constructs using either Gibson assembly or mutagenesis PCRs. All plasmids were confirmed by Sanger sequencing. Before transformation into yeast, the plasmids were digested with NotI and transformants initially selected on selective glucose medium lacking uracil and subsequently on medium containing 5-FOA to remove the URA-marker by recombination (Schrott and Osman 2023).

p426Gal1-NS-mtGFP was created by digesting p416Gal1-NS-mtGFP (Izawa et al. 2017) with SacI and EcoRI and subcloning the fragment into the SacI/ EcoRI digested pRS426 (Christianson et al. 1992).

Yeast cells were typically grown at 30°C in YP medium (10 g/L yeast extract, 20 g/L bactopeptone, pH 5.5) containing 2% of either glucose (YPD), galactose (YPGal), or glycerol (YPG), as a carbon source or in selective medium (1.7 g/L yeast nitrogen base, 5 g/L (NH_4_)_2_SO_4_, 0.02 mg/mL Adenine, 0.02 mg/mL Histidine, 0.03 mg/mL Leucine, 0.03 mg/mL Lysine, 0.004 mg/mL Tryptophan, pH 5.5) containing either 2% glucose (SD) or 2% lactate (SLac) as a carbon source.

Mitochondria were isolated from BY4742 Pth1-yEGFP and D273-10b (*Mat* α *mal*) cells grown in Lactate medium (3 g yeast extract, 1 g KH_2_PO_4_, 1 g NH_4_Cl, 0.5 g CaCl_2_x2H_2_O, 0.5 g NaCl, 1.1 g MgSO_4_x6H_2_O, 0.3 ml 1% FeCl_3_, 22 ml of 90 % lactic acid, dH_2_O to 1 L, pH 5.5, supplemented with 0.1 % glucose) at 30°C, as described in Genge et al., 2023 (Genge et al. 2023).

### Synthetic Genetic Array (SGA) analysis

SGA analysis was performed essentially as described in Tong and Boone, 2007 (A. H. Y. Tong and Boone 2007). Briefly, the MATa strain carrying query mutations (*vms1Δ::natNT2*, *ltn1Δ::hphNT1*) was constructed from the parental strain Y8205 (*MATα can1Δ::STE2pr-his5 lyp1Δ::STE3pr-LEU2 ura3Δ0 leu2Δ0 his3Δ1 met15Δ0*) (A. H. Tong et al. 2001) and mated with the deletion mutant library (Thermo Fisher) using Biomek FXP (Beckman Coulter). After selection of diploid cells, sporulation was induced by transferring cells to sporulation medium containing 1% potassium acetate. The triple deletion mutants carrying *vms1Δ::natNT2*, *ltn1Δ::hphNT1 and XXXΔ::kanMX4* were selected by transferring cells to the plates containing nourseothricin, hygromycin B and G418. The strains were then spotted onto the YPG plates, grown at 37°C for 3 days and their growth areas were measured. Each strain was generated in duplicates. Deletions with a z-score higher than 4.0 in both replicates are listed in Table S1.

### Analysis of growth of yeast cells

Growth of yeast cells was analyzed either by serial dilution spot assay on plates or in liquid media. For growth analysis on plates, yeast cells were grown in liquid YPD medium or in selective lactate medium containing 0.1% glucose ON at 30°C. ON cultures were diluted into fresh medium and grown until OD ca. 0.5. One OD of cells were then transferred into sterile tubes, cells were pelleted by centrifugation (16000 rcf, 2 min, RT) and resuspended in 1mL sterile dH_2_O. Subsequently, four ten-fold serial dilutions were made in sterile dH_2_O and 2 μl of cell suspensions were spotted on plates. The plates were incubated at indicated temperatures for 2 to 3 days.

Growth of yeast cells in liquid medium was analysed in 96-well plates using the SPECTROstar Nano (BMG Labtech). To this end, ON cultures were diluted into fresh medium and grown till OD ca. 0.5. The cells were then collected, washed three times with sterile ddH_2_O, resuspended in YPG medium to an OD_600_ of 0.1 and aliquoted into wells of a 96-well plate. Growth at 30°C was monitored every 20 minutes for up to 72h under constant shaking at 800 rpm. In every plate, each strain was grown in three separate wells and the mean of these technical triplicates was calculated in Excel. All growth curves shown represent the mean of three biological replicates, each with three technical triplicates.

### Submitochondrial localization

Isolated mitochondria were resuspended in either SH-buffer (0.6 M sorbitol, 20 mM HEPES/KOH, pH 7.4), to keep mitochondria intact, in 20 mM HEPES/KOH, pH 7.4, to burst the outer membrane and generate mitoplasts, or in 20 mM HEPES/KOH, pH 7.4 containing 0.25% Triton X-100, to solubilize mitochondrial membranes. Where indicated, proteinase K was added and the samples were incubated for 20min on ice. PMSF (1 mM) was then added to inhibit the protease and the samples were further incubated for 5 min on ice. Mitochondria and mitoplasts were reisolated by centrifugation (18000 rcf, 10 min, 4°C), resuspended in 2x Laemmli buffer (120mM Tris/HCl, pH 6.8, 20% glycerol, 4% SDS, 0.02% bromphenol blue, 3% ß-mercaptoethanol), incubated for 5 min at 95°C and then stored at -20°C until SDS-PAGE analysis. Triton X-100-solubilized sample was TCA-precipitated before resuspension in 2x Laemmli buffer and analysis by SDS-PAGE and western blot.

To analyze membrane association of proteins by carbonate extraction, isolated mitochondria were incubated in 100 mM freshly prepared Na2CO3 for 30 min on a rotating platform at 4°C. One aliquot was taken as total and TCA-precipitated. The rest of the sample was centrifuged (30 min, 186000 rcf, 2°C) and thereby separated into the supernatant fraction, which contains the soluble and peripheral membrane proteins, and the pellet fraction which contains integral membrane proteins. The pellet fraction was directly resupended in 2x Laemmli buffer whereas the supernatant was first TCA-precipitated. All samples were analyzed by SDS-PAGE and western blot.

### Protein aggregation assay

Endogenous protein aggregates were isolated essentially as previously described in Izawa et al. 2017 (Izawa et al. 2017). Briefly, yeast cells were grown in YPGal medium ON at 37°C and diluted the next day into fresh medium. When the cultures reached OD ca. 0.5, 35 OD_600_ of cells were collected by centrifugation (3220 rcf, 5min, RT), the pellets were resuspended in 500 μl IP buffer (50 mM Tris/HCl pH 7.4, 20 mM KCl, 10 mM MgCl_2_, 1mM PMSF, Protease inhibitor cocktail) and 100 μl glass beads were added. Cells were lysed by vortexing four times for 30s with 30s pauses on ice in between, the samples were diluted with 500 μl IP buffer containing 1 % Triton X-100 and incubated for 10 min at 4°C with gentle rolling. After removal of the cell debris by centrifugation (840 rcf, 10min, 2°C), an aliquot corresponding to 5 OD_600_ was taken as total (T) and the rest of the lysates were separated into soluble (Sup) and pellet (P) material by centrifugation (20000 rcf, 15min, 2°C). The pellets, containing aggregated proteins, were resolved in HU buffer (8 M urea, 5% SDS, 200 mM Tris/HCl pH 6.8, 1mM EDTA, 0.01% bromphenol blue, 2% ß-mercaptoethanol), incubated for 5 min at 95°C and stored at - 20°C until loading on SDS-PA gels. Proteins in T and Sup fractions were first TCA-precipitated (12% TCA), resolved in HU buffer, incubated for 5 min at 95°C and analyzed, together with the aggregate fraction, by SDS-PAGE followed by western blot. The material corresponding to 15 OD_600_ was loaded per lane for P and 1.5 OD_600_ per lane for T and S fractions.

### Total cell extracts

Total cell extracts for analysis of CAT-tail proteins were prepared essentially as described in Izawa et al. 2017 (Izawa et al. 2017). Briefly, expression of NS-mtGFP from the p426*GAL1*-NS- mtGFP plasmid was induced by addition of 0.5% galactose for 20h to logarithmically growing cells in selective lactate medium at 30°C. Cells (3 OD_600_) were harvested by centrifugation (5000 rcf, 5 min, RT) and resuspended in 1 mL ice-cold H_2_O before 150 μl of solution containing 2 M NaOH and 7 % ß-mercaptoethanol were added. After 10 min incubation on ice, TCA was added to 12% final concentration and samples were incubated for an additional 30 min on ice. The precipitated proteins were isolated by centrifugation (17000 rcf, 10min, 2°C), washed with 1 mL ice-cold acetone and the final pellet was resuspended in 80 μl HU-buffer, incubated for 5 min at 95°C and analyzed by SDS-PAGE (1.5 OD_600_ per lane) followed by western blot.

### Peptidyl-tRNA accumulation assay

To assess the presence of peptidyl-tRNAs in cells, the protocol described in Verma et al. 2013 (Verma et al. 2013) was modified. Briefly, yeast cells (5 OD_600_) were washed three times with 1 mL ice cold buffer containing 50 mM Tris/HCl, pH7.5, 10 mM NaN_3_, 10 mM EDTA, 10 mM EGTA, 1x protease inhibitor tablet (Roche), 10 mM NEM, 50 mM NaF, 60 mM ß-glycerophosphate, 10 mM sodium pyrophosphate, and split into two 200 μl aliquots. The cells were reisolated and 50 μl glass beads were added together with 40 μl SDS buffer (120 mM MOPS/NaOH, pH 6.8, 20 % glycerol, 4 % SDS, 0.02 % bromphenol blue) containing either 1 % DEPC and 2 % ß-mercaptoethanol or 200 ng/µl RNaseA. Cells were incubated at 65°C for 5 min and vigorously vortexed for 1 min. ß-mercaptoethanol was added to 2% to RNaseA-treated samples. All samples were incubated for 3 min at 95°C, centrifuged for 1 min at 16000 rcf and directly loaded on 4-12 % NuPAGE gels (Thermo fisher). Gels were run in MOPS buffer (Thermo fisher) according to manufacturer’s instructions and subsequently blotted on nitrocellulose membranes.

### Immunoprecipitation of NS-mtGFP

Expression of NS-mtGFP was induced in yeast cells for 20h, as described above. Cells (180 OD_600_) were collected by centrifugation (3220 rcf, 5min, RT) and resuspended in 600 μl lysis buffer (25 mM Tris/HCl, pH 7.5, 50 mM KCl, 10 mM MgCl_2_, 1 mM EDTA, 5% glycerol, 1x protease inhibitor tablet (Roche)). Glass beads (50 μl) were added and the cells were vortexed four times for 30s with 30s pauses on ice. The samples were diluted with 600 μl lysis buffer containing 1% Triton X-100 and incubated for 10 min on a rolling platform at 4°C. After a clarifying spin (1000 rcf, 10 min, 4°C), 1 mL of lysate was added to 10 μl GFP-Trap^®^ Magnetic Particles M-270 (Chromotek/ Proteintech) and incubated for 2h at 4°C on a rotating platform. During this time, the remaining lysate was diluted with ice cold water to 1 mL and the proteins therein TCA-precipitated (12% TCA). GFP-Trap beads were isolated using a magnetic rack, the non-bound fraction was removed and the beads were subsequently washed three times for 10 min with 500 μl lysis buffer containing 0.05% Triton X-100. The specifically bound proteins were either directly eluted with HU buffer and incubation for 5 min at 95°C and analyzed by SDS-PAGE and western blot or were directly used for on bead digest and mass spectrometry. For mass spectrometry, three biological replicates of each strain were analyzed. The beads were washed twice with 50 mM Tris/HCl, pH 7.5, and subsequently with 50 mM Tris/HCl, pH 7.5 containing 2 M urea. For the first trypsin digest, the beads were resuspended in 160 µl of 50 mM Tris/HCl, pH 7.5, 1 M urea, 1 mM Dithiothreitol (DTT), and 0.8 µg of trypsin (Pierce, Thermo Scientific) were added and incubated for 3 h at 25°C with continuous shaking. Post-incubation, the supernatant was saved and the beads were washed twice with 120 µl 50 mM Tris/HCl, pH 7.5 containing 1 M urea. The supernatants were combined with the trypsin digest. To the pooled sample, DTT was added to a final concentration of 4 mM, and the sample was incubated at 25°C for 30 min with shaking. Iodoacetamide was added to a final concentration of 10 mM and the sample was incubated at 25°C for 45 min with shaking in the dark. Subsequently, an additional 1 µg of trypsin was added, and the second digestion was allowed to proceed overnight at 25°C with continuous shaking. The combined digested peptides were purified using home-made C18 stage tips (Rappsilber, Ishihama, and Mann 2003).

### LC-MS/MS analysis

Liquid chromatography tandem mass spectrometry (LC-MS/MS) was performed on a nano-LC system (Ultimate 3000) coupled to an Impact II Q-TOF (Bruker Daltonics, Bremen, Germany) using a CaptiveSpray nano electrospray ionization (ESI) source as described previously (Espinoza-Corral, Schwenkert, and Schneider 2023). Peptides (1 µg) were separated over a 30 min linear gradient of 5–45% (v/v) acetonitrile on a Acclaim Pepmap RSLC analytical column (C18, 100 Å, 75 µm x 50 cm) (Thermo Scientific). MS1 spectra with a mass range of m/z 200– 2000 were acquired at 3 Hz and the 18 most intense peaks were selected for MS/MS analysis with an intensity-dependent spectrum acquisition rate between 4 and 16 Hz. Dynamic exclusion duration was set to 0.5 minutes.

Raw data files were analyzed using MaxQuant software version 2.4.4.0 (Cox and Mann 2008), with peak lists compared against the Saccharomyces cerevisiae reference proteome from UniProt (www.uniprot.org). All settings were kept at default values and protein quantification was performed using the label-free quantification (LFQ) algorithm (Cox et al. 2014). Further analysis was performed using Perseus version 2.0.9.0 (Tyanova, Temu, and Cox 2016). Potential contaminants, proteins identified only through site modification, and reverse hits were removed. Only proteins quantifiable by the LFQ algorithm in at least two out of three replicates were kept. LFQ intensities were transformed to log_2_ values and missing data were imputed from a normal distribution within Perseus using standard parameters. Samples were statistically compared using a t-test and data was visualized using a custom RStudio (RStudio Team 2019) script.

### Fluorescence microscopy

Yeast cells were grown in selective lactate medium containing 0.1% glucose ON at 30°C. On the next day, the cells were diluted into fresh selective lactate medium without any added sugar and grown till OD ca. 0.5. One sample per culture was directly imaged (0h) and the rest of the cultures were diluted to OD_600_ of 0.3 and expression of NS-mtGFP was induced by addition of galactose to 0.5%. If needed, the cultures were diluted in selective lactate medium containing 0.5% galactose so that OD_600_ remained between 0.3 and 0.8. After indicated time periods, 0.15 OD_600_ were isolated by centrifugation (3000 rcf, 3 min, RT), resuspended in 1 mL staining buffer (10 mM Hepes, pH 7.6, 5% glucose, sterile filtered) containing 50 nM Image-iT TMRM-Reagent (Thermo fisher) and rotated for 15 min at 30°C in the dark. Cells were re-isolated by centrifugation and washed two times with 1 mL staining buffer and once with 1 mL selective lactate medium containing 0.5% galactose. Cells were finally resuspended in 500 μl medium and 300 μl of the cell suspension were transferred into Concanavalin A coated 8- well chambered μ-slides with glass bottom (Ibidi). The cells were attached to Con A by a 2 min centrifugation of the slides at 800 rcf. Non-bound cells were subsequently washed three times with 400 μl medium. Images were captured in Z-stacks (41 slices, 200 nm) with a Nikon ECLIPSE Ti2-E microscope (Nikon MEA54000), with a CFI Apochromat TIRF 100XC oil immersion objective (Nikon MRD01991) and a Photometrics Prime 95B 25-mm camera (PH-P95B-25) at 30°C. The ND2 files of the NIS-ELEMENTS AR software (Nikon MQS31000) were converted to OME-TIFF using a custom Python 3 macro and deconvolved using Huygens Essential v 18.10 (Scientific Volume Imaging). Pictures in figures are presented as maximum intensity Z-projections created with Fiji/ImageJ. To allow for better visualization in case of the GFP channel pictures were equally and in case of the TMRM signal individually adjusted by setting the upper and lower borders of depicted pixel intensities.

### Image quantification

Single cells expressing GFP were manually cropped out using the Fiji ROI function and quantified with a custom Fiji/ImageJ macro. In short, the GFP channel of each cropped image was automatically analyzed for particles with the Fiji plugin “3D Objects Counter” (“slice=20 min.=7 max.=2562500 exclude_objects_on_edges objects surfaces statistics summary”) (Bolte and Cordelières 2006). Initially, images were manually analyzed and evaluated to identify errors in the macro’s performance. Subsequently, these erroneous analyses were automatically excluded by the macro using a predefined threshold. The data were then analyzed and plotted using custom Python scripts. Average values of three independent experiments with at least 150 cells per replicate were statistically compared using t-test, as recommended by Lord et al., 2020 (Lord et al. 2020).

