## Supplementary Figures for "Delayed protein translocation protects mitochondria against toxic CAT-tailed proteins"

**Supplementary Figure 1: Pth2 is peptidyl-tRNA hydrolase involved in the yeast mitoRQC**

[A, C] Growth of indicated yeast deletion strains was analyzed by a serial dilution assay. Serial dilutions of cells in logarithmic phase were spotted on plates containing glucose as a fermentable (YPD) or glycerol (YPG) as a non-fermentable carbon source. The plates were incubated at 30°C or 37°C.

[B, G, I] Indicated strains were grown to log phase in YPD, washed and diluted to an $OD_{600}$ of 0.1 in YPG. Growth was monitored under constant shaking at 30°C.

[D] Alignment of Pth2 homologues from *Saccharomyces cerevisiae* (Sc_Pth2), *Schizosaccharomyces pombe* (Sp_Pth2), *Drosophila melanogaster* (Dm_CG1307), *Danio rerio* (Dr_Ptrh2) and *Homo sapiens* (Hs_Ptrh2). Based on the degree of primary sequence conservation, Pth2 is divided in an N- (1-84) and a C-terminal (85-208) region. Predicted transmembrane domains (TMHMM-server) in the N-terminal regions of all homologues are indicated with red rectangles. The conserved Pth2 active site is marked in yellow. Asterisk indicates the aspartic acid residue D174 that is essential for the catalytic activity of Pth2.

[E] Pth2 amino acid sequence was analyzed for the presence of transmembrane domains using TMHMM prediction server. The server predicted one transmembrane domain (residues 7-29).

[F] Schematic illustrations of Pth2 constructs used in the study. Pth2 is divided in N- and C-terminal regions. Pth2N-GFP harbours residues 1-84 of Pth2 followed by full length superfolder GFP (1-238). $TM_{Tom70}$-Pth2 consists of the first 40 residues of Tom70, which were previously shown to target and insert passenger proteins into the mitochondrial outer membrane, fused to residues 34-208 of Pth2. $Pth2^{D174N}$ has a point mutation changing aspartic acid residue at position 174 to an asparagine residue rendering Pth2 catalytically deficient as a peptidyl-tRNA hydrolase. $Pth2^{\Delta TM}$ lacks residues 1-33 of Pth2 creating a non-membrane bound form of Pth2 without affecting its catalytic activity. Pth2N-Pth1 contains residues 1-84 of Pth2 fused to residues 7-191 of Pth1 (residues 1 to 6 of Pth1 contain a predicted N-terminal mitochondrial matrix targeting signal using MitoFates 1.2). Pth2N-$Pth1^{H24N}$ additionally contains a point mutation where the histidine residue at position 24 in the original Pth1 sequence was mutated to an asparagine residue (position 101 in the fusion construct). This mutation in human Ptrh1 was previously published to render the enzyme catalytically deficient [32].

[H] Alignments of Pth1 homologues from *Saccharomyces cerevisiae* (Sc_Pth1), *Schizosaccharomyces pombe* (Sp_Pth1), *Danio rerio* (Dr_Ptrh1) and *Homo sapiens* (Hs_Ptrh1).

Asterisk indicates the conserved histidine residue essential for the Ptrh1 catalytic activity.

*ΔΔ* = *Δvms1 Δltn1*; *ΔΔΔ* = *Δvms1 Δltn1 Δpth2*

**Supplementary Figure 2: Fluorescence microscopy of NS-mtGFP expressing cells**

[A] Related to Fig 2C. Fluorescence microscopy of NS-mtGFP expressing cells was done 20h after induction of NS-mtGFP expression. Intensity of the GFP channel was adjusted so that signals in WT and *ΔΔΔ* are visible as well. In contrast to *ΔΔ* and *ΔΔΔ* cells which primarily contain GFP signals in aggregates, WT and *ΔΔΔ* show typical mitochondrial staining.

*ΔΔ* = *Δvms1Δltn1*; *ΔΔΔ* = *Δvms1Δltn1Δpth2*; *ΔΔΔ* = *Δvms1Δltn1Δrqc2*

**Supplementary Figure 3: Expression of mitoRQC substrates is toxic in cells lacking Rqc2**

[A] Related to 2B and 3B. Growth of cells expressing NS-mtGFP. Indicated yeast cells were grown overnight in selective lactate medium containing 0.1% glucose at 30°C. The next day, the cultures were diluted in selective lactate medium without any additional sugar and grown till $OD_{600}$ 0.4-0.6. Serial dilutions of cells were prepared and spotted on plates containing either 0.5% galactose to induce expression of NS-mtGFP or 0.1% glucose, as a control. Plates were incubated at indicated temperatures.

*ΔΔ* = *Δvms1Δltn1*; *ΔΔΔ* = *Δvms1Δltn1Δpth2*; *ΔΔΔ* = *Δvms1Δltn1Δrqc2*

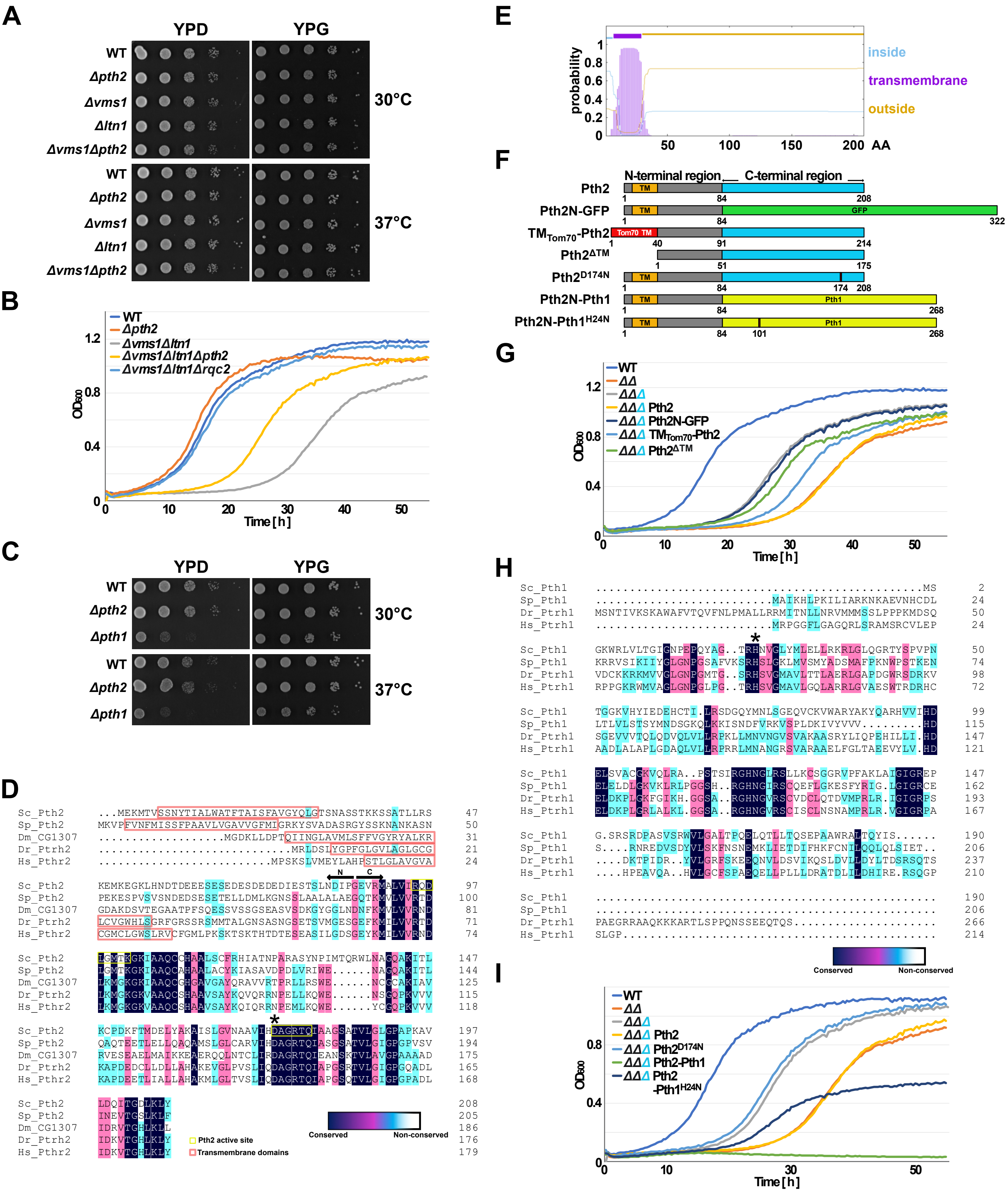


**Figure S1: Bertram et al.**

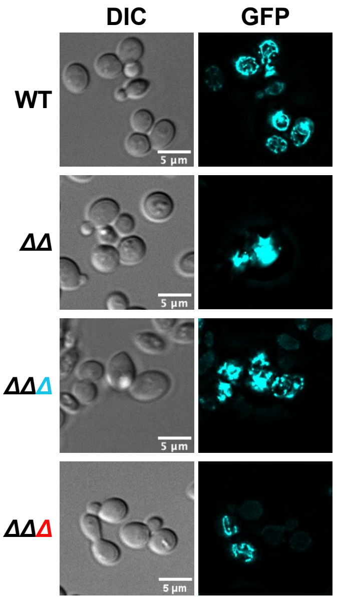


**Figure S2: Bertram et al.**

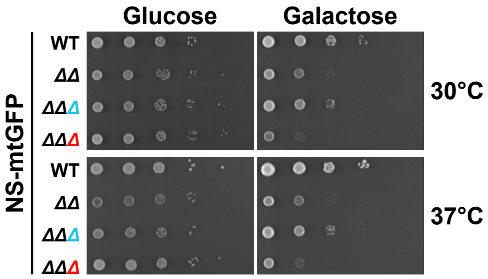


Figure S3: Bertram et al.
